# Combinatorial Modulation of Wnt, STAT3, TGF-β, and Tie2 Pathways Drives Brain Endothelial Cell–Like Differentiation from hiPSCs

**DOI:** 10.64898/2026.08.21.746025

**Authors:** Jung Hyun Lee, Erin S. O’Connor, Ji Ye Lee, Kristina M. Holton, Lee L. Rubin

## Abstract

During development, endothelial cells (ECs) migrate into the brain and acquire blood–brain barrier (BBB) properties such as tight junctions, limited transcellular transport, and high electrical resistance. Although key signaling pathways that are active in vivo have been identified, factors critical in inducing brain EC differentiation *in vitro* remain unclear. Here, we describe conditions that promote brain EC–like gene expression in human pluripotent stem cell (hiPSC)–derived ECs. Activation of Wnt/β-catenin signaling upregulates the brain EC marker GLUT1 (SLC2A1) while suppressing the peripheral EC marker PLVAP. Simultaneously, stimulation of STAT3 by CNTF together with TGF-β inhibition increases CLDN5 expression. We further found that hiPSC-derived ECs secrete high levels of angiopoietin-2 (ANGPT2) and that razuprotafib (AKB-9778), a PTPRB (VE-PTP) inhibitor, inhibits ANGPT2 and improves monolayer integrity. These results suggest that combinatorial modulation of specific signaling pathways stimulates the differentiation of human brain ECs *in vitro*.

**Highlights:**

- Wnt signaling induces GLUT1 (SLC2A1) and reduces PLVAP expression in hiPSC-derived ECs.
- CNTF addition and inhibition of TGFBR1 upregulate CLDN5.
- Activation of Tie2 signaling enhances barrier function.
- Combining these factors induces brain EC-like phenotypes in hiPSC-derived ECs.

## Introduction

The blood-brain barrier (BBB) is a highly selective barrier formed by endothelial cells (ECs), pericytes, astrocytes, and vascular smooth muscle cells, and maintains the homeostasis of the central nervous system (CNS)(Engelhardt, 2003; Langen et al., 2019; Obermeier et al., 2013). Brain ECs exhibit unique characteristics such as complex high resistance tight junctions, a low rate of transcytosis, and polarized transporter expression which distinguish them from peripheral ECs and combine to constitute the BBB. In addition to their role as a protective barrier, the brain blood vessels dynamically regulate blood flow in response to changes in neural activity (Friedman et al., 2025; Iadecola, 2017) and are known to secrete factors that regulate CNS neurogenesis (Karakatsani et al., 2019; Shen et al., 2004; Sosa et al., 2007). More recent studies have shown that brain ECs serve a key role in regulating to brain’s response to circulating blood factors, acting in some cases as the target and in others, by controlling their entry into the brain (Chen et al., 2020; Katsimpardi et al., 2014; Ximerakis et al., 2023; Yang et al., 2020). Importantly, BBB dysfunction is often accompanied by aging and neurodegenerative diseases, suggesting a critical association between brain ECs and brain function.

The BBB is formed through a complex multi-stage process. During development, ECs outside the brain invade the neural tissue and differentiate into the highly specialized brain ECs that constitute the BBB(Bautch and James, 2009; James and Mukouyama, 2011; Segarra et al., 2019). Transplant studies using chick and quail ECs suggest that they are highly plastic and able to acquire brain specific characteristics under the influence of local cues(Stewart and Wiley, 1981). VEGF derived from neural tissue is known to guide vessel sprouts into the brain parenchyma and influence EC permeability (Haigh et al., 2003; Hogan et al., 2004; James et al., 2009). In mice, studies using genetic ablation of Wnt signaling components like Wnt7a, Wnt7b, and β−Catenin showed that Wnt signaling plays essential roles in CNS vascularization as well as the acquisition of BBB properties during brain development(Daneman et al., 2009; Liebner et al., 2008; Stenman et al., 2008). In addition to the Wnt pathway, studies have discovered that pathways regulated by Hedgehog (Alvarez et al., 2011; Wang et al., 2021a), retinoic acid(Mizee et al., 2013), and TGF-β (Nguyen et al., 2011; Schumacher et al., 2023) play roles in the development or maintenance of the BBB.

Despite an extensive number of studies using animal models, not much is known regarding which signaling pathways are sufficient to induce or maintain brain EC differentiation in vitro. This may reflect the extreme plasticity of ECs. For instance, studies on isolated primary brain ECS show that maintaining barrier properties outside the brain is challenging. Primary brain ECs isolated from animal models or immortalized cell lines derived from human brain microvessels do not fully express brain EC-enriched genes (Calabria and Shusta, 2008; Helms et al., 2015a; Lyck et al., 2009; Sabbagh and Nathans, 2020a; Steiner et al., 2011; Urich et al., 2012). Wnt/β-catenin signaling is essential during *in vivo* BBB development, and its activation induces partial conversion of leaky vessels in circumventricular organs to barrier-forming vessels by reducing levels of PLVAP and increasing expression of the tight junction protein CLDN5 (Wang et al., 2019). However, Wnt/β-catenin activation in isolated primary mouse brain ECs does not fully maintain downstream brain EC gene expression(Sabbagh and Nathans, 2020a). This suggests that other pathways active *in vivo* also need to be regulated *in vitro*.

Human *in vitro* BBB models range from immortalized brain EC lines like hCMEC/D3 to multicellular microfluidic systems(He et al., 2014; Helms et al., 2015b). More recently, human induced pluripotent stem cells (hiPSCs) have been used to establish brain EC models using a variety of approaches including small molecule treatment (Praça et al., 2019; Roudnicky et al., 2020a), transcriptional reprogramming (Cui et al., 2025; Lu et al., 2021; Roudnicky et al., 2020b), and coculture with other BBB cells such as pericytes and astrocytes(Nahon et al., 2024; Nishihara et al., 2020; Praça et al., 2019). However, several limitations remain. Most models exhibit only modest transendothelial electrical resistance (TEER), a measure of paracellular permeability, along with variable expression of brain EC genes. Conversely, some iPSC-derived BBB models that achieved extremely high TEER exhibit epithelial cell markers with lower expression of EC markers(Lippmann et al., 2012; Lu et al., 2021; Qian et al., 2019). Taken together, continued development of differentiation protocols and understanding of molecular pathways to induce brain EC differentiation is needed for studying the physiological roles of ECs in development, aging, and neurodegenerative disease.

Here, we demonstrate the converging roles of multiple signaling pathways to induce brain EC differentiation of hiPSCs. We adopted a stepwise approach, first generating ECs from hiPSCs and then differentiating the ECs into more brain-like ECs that exhibit BBB properties. Combined modulation of Wnt, STAT3, and TGF-β signaling regulates expression of a subset of brain EC genes and improves barrier function. In addition, we discovered that hiPSC-derived ECs express high levels of angiopoietin 2 (ANGPT2), a factor that increases vascular permeability, and counteracting high ANGPT2 pathway significantly improves barrier integrity. Our study shows that a multi-faceted approach targeting multiple signaling pathways is needed for *in vitro* brain EC differentiation and provides critical knowledge to develop a functional *in vitro* BBB model.

## Results

### Canonical Wnt signaling activation upregulates GLUT1 and downregulates PLVAP in ECs

First, we characterized the endothelial differentiation process from hiPSCs by measuring the expression of key lineage marker genes throughout EC differentiation. We adapted a published protocol that involves mesoderm induction, followed by endothelial specification, magnetic sorting, and expansion of purified ECs (Patsch et al., 2015a). To validate the differentiation trajectory, we measured mRNA levels of marker genes at various times (Fig S1A). Upon mesoderm induction with CHIR99021 and BMP4, expression of the pluripotency gene NANOG decreased, while the mesoderm markers BRYCHYURY and PAX2 were robustly induced. On day 4, mesodermal lineage cells were exposed to VEGF and forskolin, which promoted endothelial specification. Correspondingly, VEGF receptor genes FLT1 and KDR, and pan-endothelial markers PECAM1 (CD31), CDH5 (CD144; VE-Cad), and SOX17 were strongly induced and remained high thereafter. The resulting EC population, purified by magnetic sorting using CD144 microbeads, was highly homogeneous for CD31 and CDH5 (Fig S1B) and did not express ectodermal markers, PAX6 and MAP2, or epithelial lineage markers, CDH1 and EPCAM, confirming successful EC differentiation.

To assess the role of Wnt signaling in brain EC differentiation, we examined changes in the expression of selected genes including: GLUT1 (SLC2A1), a major glucose transporter highly expressed in brain ECs; CLDN5, which encodes an essential tight junction protein; PLVAP, a marker of peripheral and fenestrated endothelium; and MFSD2A, a lipid transporter that regulates transcytosis. hiPSC-derived ECs were treated with the Wnt ligands WNT3A, WNT5A, WNT5B, WNT7A, as well as the GSK3b inhibitor CHIR99021. Canonical Wnt activators WNT3A and CHIR99021 significantly upregulated GLUT1 and APCDD1 (a known target of canonical Wnt signaling), while simultaneously downregulating PLVAP (Fig 1A and S2). Consistent with the qPCR data, immunostaining for GLUT1 and PLVAP confirmed the effects of Wnt activation (Fig 1B and 1C). Imaging of GFP-tagged β-Catenin revealed that a high concentration of CHIR99021 promoted nuclear accumulation of β-Catenin, whereas WNT3A treatment did not significantly alter nuclear β-Catenin levels. This indicates that GLUT1 induction can occur independently of an increase of nuclear β-Catenin. In contrast, other prominent brain EC genes like CLDN5 and MFSD2A were not significantly regulated by Wnt activation, suggesting that additional signaling pathways contribute to brain EC differentiation *in vitro*.

**Figure 1.**
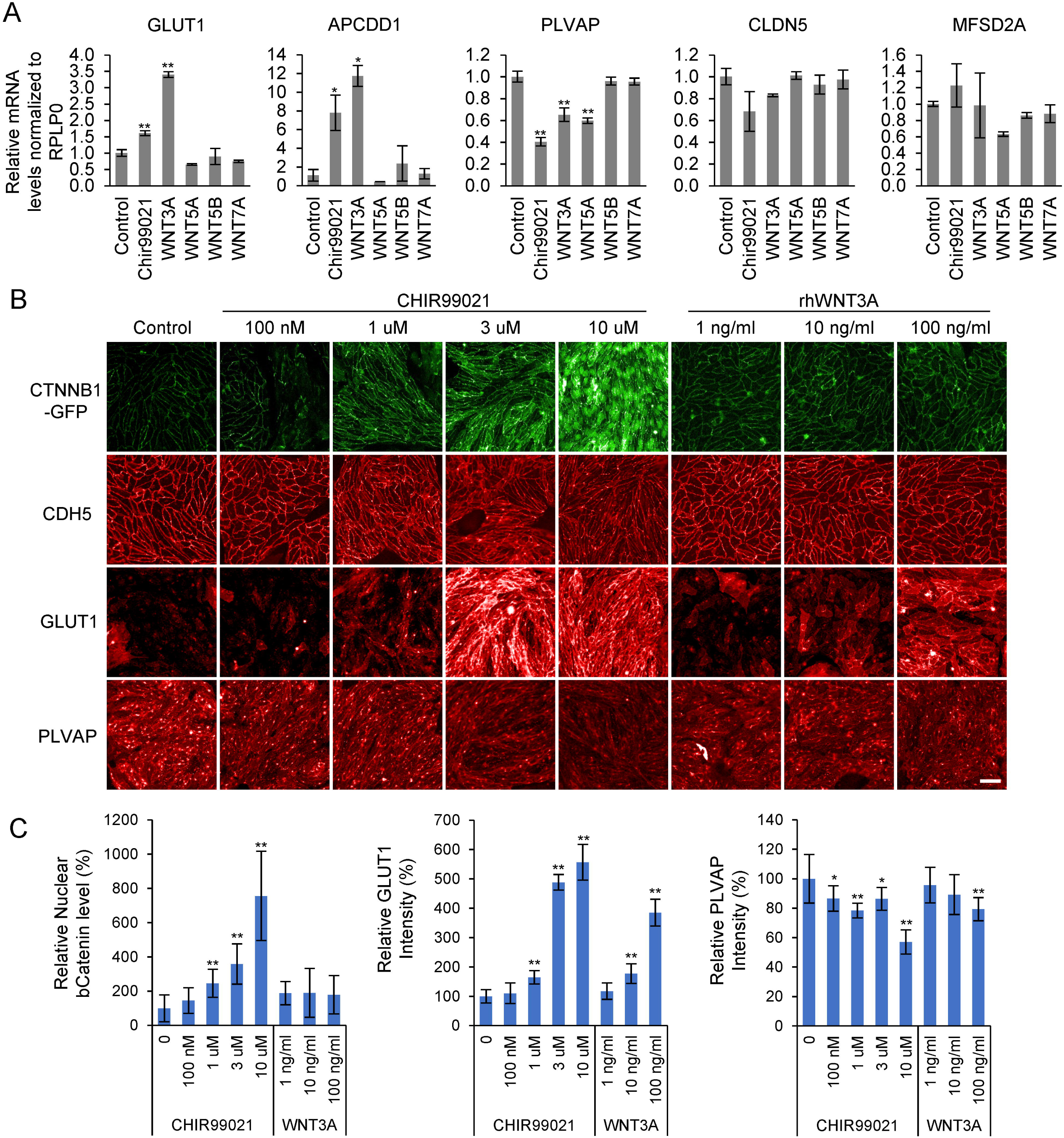
WNT activation upregulates GLUT1 and downregulates PLVAP in hiPSC-derived ECs. (A) mRNA expression of brain EC-associated genes in hiPSC-derived ECs after 2 days of treatment with WNT ligands (100 ng/ml) or the GSK3β inhibitor Chir99021 (1 uM). (B) Representative images of GFP-tagged CTNNB1 (β-catenin), CDH5 (VE-cadherin; CD144), GLUT1, and PLVAP in hiPSC-ECs after 2 days of treatment with varying concentrations of Chir99021 or WNT3A. The CTNNB1::GFP iPSC line (Allen Cell, AICS-0058-067iPSC) was used to visualize β-catenin distribution. Immunostaining was performed for CDH5, GLUT1, and PLVAP. (C) Quantification of nuclear β-catenin intensity, total GLUT1, and total PLVAP expression. Data are represented as mean ± standard deviation. Scale bar, 50 um. *, P<0.05 vs. Control; **, P<0.01 vs. Control.

We additionally examined other signaling pathways implicated in BBB development, including retinoic acid (RA) and Sonic hedgehog (SHH) signaling. RA decreased GLUT1 and PLVAP, but did not significantly alter CLDN5 or MFSD2A. Activation of the Hedgehog pathway with recombinant Shh, purmorphamine, or Smoothened agonist (SAg) did not significantly affect the expression of GLUT1, CLDN5, PLVAP, or MFSD2A (Fig S3). Thus, neither RA nor Hedgehog pathway activation promoted a consistent brain EC–like gene expression pattern under the conditions tested.

### The CNTF and STAT3 pathway increased CLDN5 expression

Next, we examined the effects of neurotrophic factors present in the developing brain on brain EC differentiation. We tested glial cell-line derived neurotrophic factor (GDNF), brain-derived neurotrophic factor (BDNF), and ciliary neurotrophic factor (CNTF). Of those factors, only CNTF increased the expression of CLDN5 (Fig 2A, 2B, and S4). CNTF is a member of the interleukin 6 (IL-6) family cytokines and is produced by various glial cells in the central nervous system. CNTF acts through a receptor complex composed of interleukin 6 cytokine family signal transducer (IL6ST; gp130) and CNTF receptor alpha (CNTFRa) (Dallner et al., 2002; Rose-John, 2018; Stöckli et al., 1991). The CNTF effect was higher with co-treatment with a soluble form of CNTF receptor alpha (CNTFRa) (Fig 2A and 2B). Furthermore, treatment with CNTF and CNTFRa increased GLUT1 and CLDN5 mRNA expression whereas PLVAP and MFSD2A expression was not significantly changed (Fig 2C). Consistent with previous studies(Hirano et al., 1994; Rapp et al., 2023), CNTF induced phosphorylation of STAT3 (signal transducer and activator of transcription 3), which persisted for at least 2 days after stimulation (Fig 2D). To further verify the role of STAT3 in CLDN5 expression, we treated cells with the JAK inhibitor ruxolitinib and the STAT3 inhibitor C188-9 in the presence of CNTF and CNTFRa. Both ruxolitinib and C188-9 reduced the CNTF-mediated increase in CLDN5 (Fig 2E and 2F). Consistently, western blot analysis showed that the CNTF/CNTFRa induced CLDN5 upregulation was eliminated by C188-9, which effectively reduced STAT3 phosphorylation (Fig 2G). Among other IL-6 family cytokines, Oncostatin M (OSM) also increased CLDN5 expression whereas IL-6 and leukemia inhibitory factor (LIF) did not (Fig S5). These differences are likely due to the availability of their respective receptors (e.g., IL-6R, OSMR, LIFR) on hiPSC-derived ECs. Together, these data suggest that CNTF, in the presence of CNTFRa, acts as an inducer of CLDN5 expression in ECs via STAT3 signaling. The effect of CNTF signaling on GLUT1 appears weaker than that of WNT activation, while CNTF-STAT3 signaling has minimal impact on PLVAP, and MFSD2A expression.

**Figure 2.**
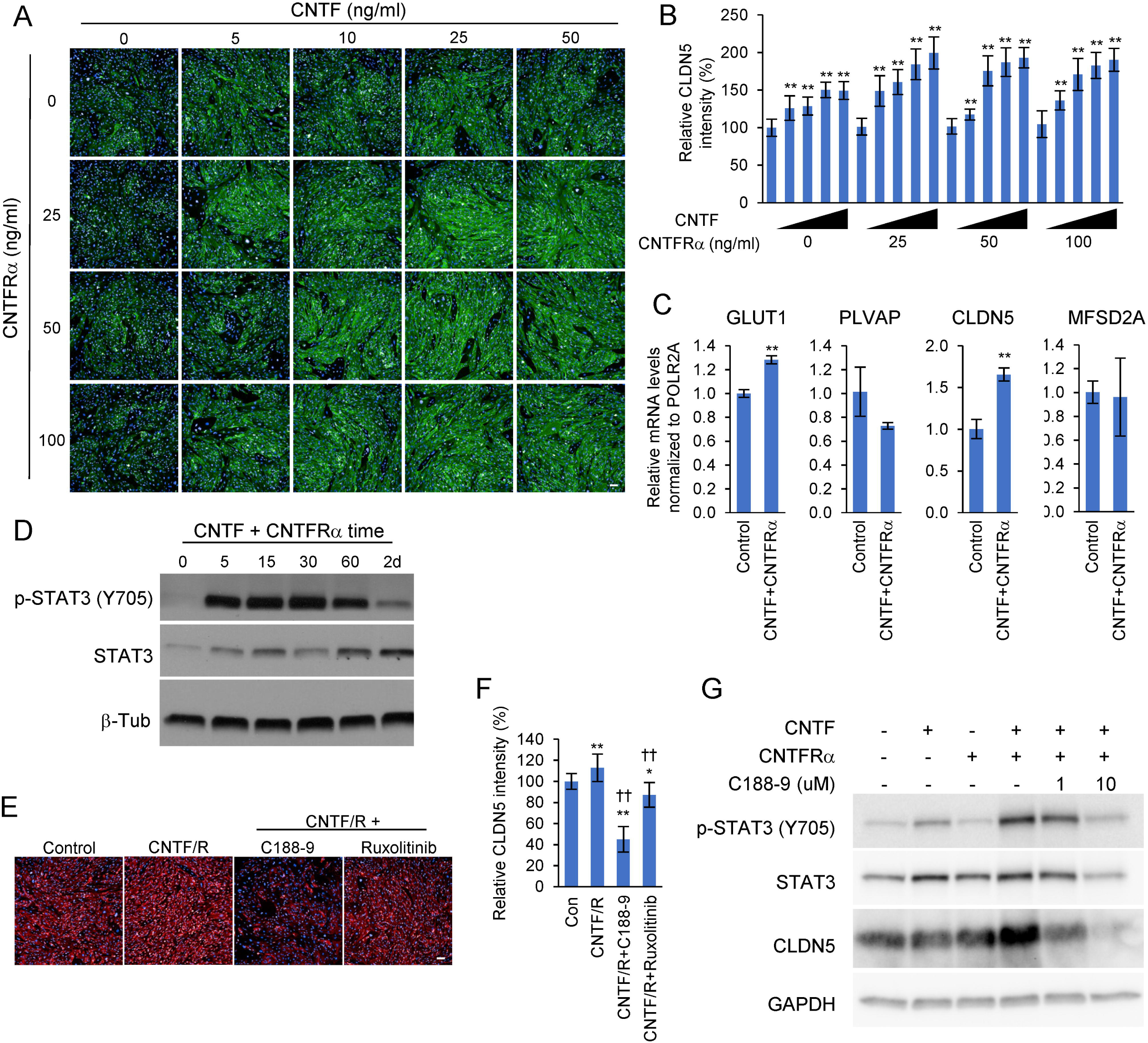
CNTF-STAT3 pathway upregulates CLDN5 in hiPSC-derived ECs. (A and B) Representative images and quantification of hiPSC-derived ECs treated with increasing doses of recombinant human CNTF and CNTFRα. (C) Relative mRNA expression of brain EC-associated genes after 2 days of treatment with CNTF (100 ng/ml) and CNTFRα (50 ng/ml). (D) Western blot showing STAT3 phosphorylation after CNTF and CNTFRα treatment. (E and F) Representative images and quantification of CLDN5 immunostaining in cells treated with CNTF and CNTFRα in the presence or absence of C188-9 (10 uM) or Ruxolitinib (10 uM). (G) Western blot showing STAT3 phosphorylation and CLDN5 protein levels in hiPSC-derived ECs treated with CNTF, CNTFRα, and/or C188-9 for 2 days. Data are represented as mean ± standard deviation. Scale bars, 100 um. *, P<0.05 vs Control; **, P<0.01 vs Control; ††, P<0.01 vs CNTF/CNTFRα.

### TGF-**β** pathway inhibition increases CLDN5 expression

To evaluate the role of the TGF-β pathway in brain EC differentiation, we treated hiPSC-derived ECs with multiple TGF-β receptor (TGFBR1; ALK5) inhibitors: RepSox, SB431542, SB525334, and Galunisertib. All four inhibitors increased CLDN5 expression (Fig 3A and 3B), consistent with previous studies(Roudnicky et al., 2020a; Watabe et al., 2003). Among them, RepSox was the strongest inducer of CLDN5. Notably, RepSox also upregulated GLUT1 and downregulated PLVAP expression suggesting that TGF-β pathway inhibition may promote brain EC differentiation more broadly. However, no significant changes in MFSD2A expression were observed. qPCR analysis further confirmed that RepSox increased CLDN5 and GLUT1 mRNA levels and suppressed PLVAP, while expression of MFSD2A remained unchanged (Fig 3C). Finally, co-treatment with RepSox and CNTF/CNTFRa resulted in an additive increase in CLDN5 level, indicating that these two pathways act independently to regulate CLDN5 expression (Fig 3D and 3E).

**Figure 3.**
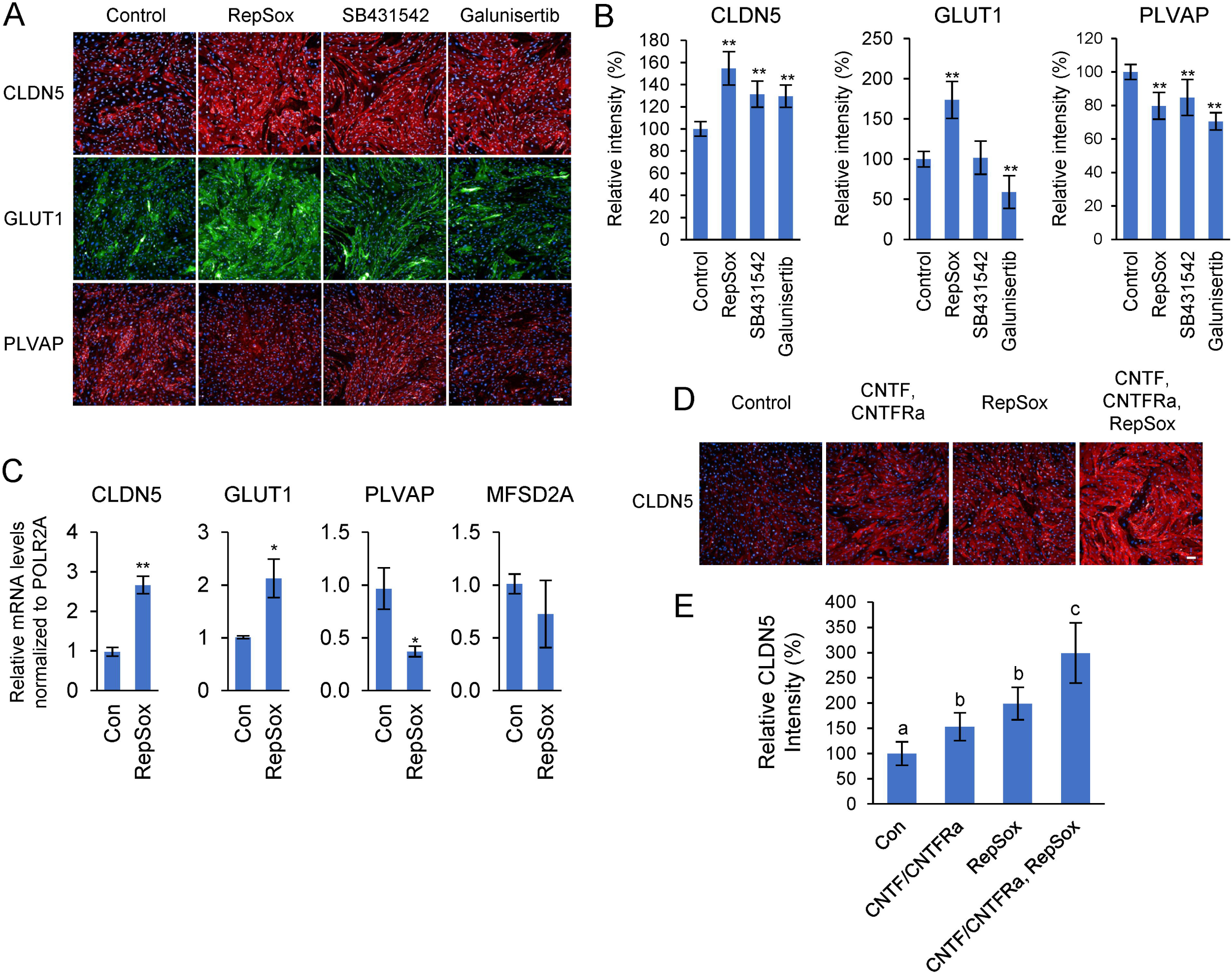
Inhibition of TGFBR1 upregulates CLDN5 in hiPSC-derived ECs. (A and B) Representative images and quantification of hiPSC-ECs treated with different TGFBR1 inhibitors (10 uM, 2 days). (C) Relative mRNA expression of CLDN5, GLUT1, PLVAP, and MFSD2A in hiPSC-ECs treated with 10 uM RepSox for 2 days. (D) Immunostaining for CLDN5 in cells treated with CNTF/CNTFRa, RepSox or both for 2 days. CNTF, 25 ng/ml; CNTFRa, 50 ng/ml; RepSox, 10 uM. (E) Quantification of CLDN5 staining. One-way ANOVA with Tukey’s HSD was used for multiple comparisons. Different letters indicate statistically significant differences (P<0.05). Data are represented as mean ± standard deviation. Scale bars, 100 um., *, P<0.05 vs Control; **, P<0.01 vs Control in t-test.

### Combinatorial modulation of Wnt, CNTF-STAT3, and TGF-**β** pathways promotes the transcriptional program of brain EC differentiation

We next tested whether a combination of WNT3A, CNTF, CNTFRa, and RepSox (hereafter referred to as WCNR) could induce broader brain EC-like transcriptional changes. To be sure that the changes we observed were not limited to a single iPSC line, we tested the effect of WCNR in multiple EC lines in addition to DiPS-1016-SevA derived ECs. When WCNR was added to ECs derived from DiPSC-1016-SevA, BJ SiPS, WTC-mEGFP-CTNNB1, and PGP-1, all four iPSC-EC lines exhibited increased GLUT1, APCDD1, and CLDN5 and decreased PLVAP mRNA levels (Fig 4A). Also, we observed a small but significant increase of MFSD2A, suggesting that combinatorial treatment is promoting BBB-like gene expression profiles. In two primary human EC lines HUVEC (Lonza) and hBMVEC (iXCell Biotechnologies), WCNR increased expressions of CLDN5, APCDD1, and decreased PLVAP. However, GLUT1 was not induced in either HUVEC or hBMVEC by WCNR, implying that signaling pathways required to induce GLUT1 in these primary ECs might be different from iPSC-derived ECs. Correspondinly, immunostaining and western blot analysis also showed increased CLDN5 in all tested ECs but increased GLUT1 only in iPSC-derived ECs but not in primary ECs. (Fig 4B, S6, and 4C). WCNR treatment also produced a significant increase in barrier integrity in most cell lines, as measured by transendothelial electrical resistance (TEER) (Fig 4D), indicating that TEER is closely associated with levels of tight junction proteins like CLDN5.

**Figure 4.**
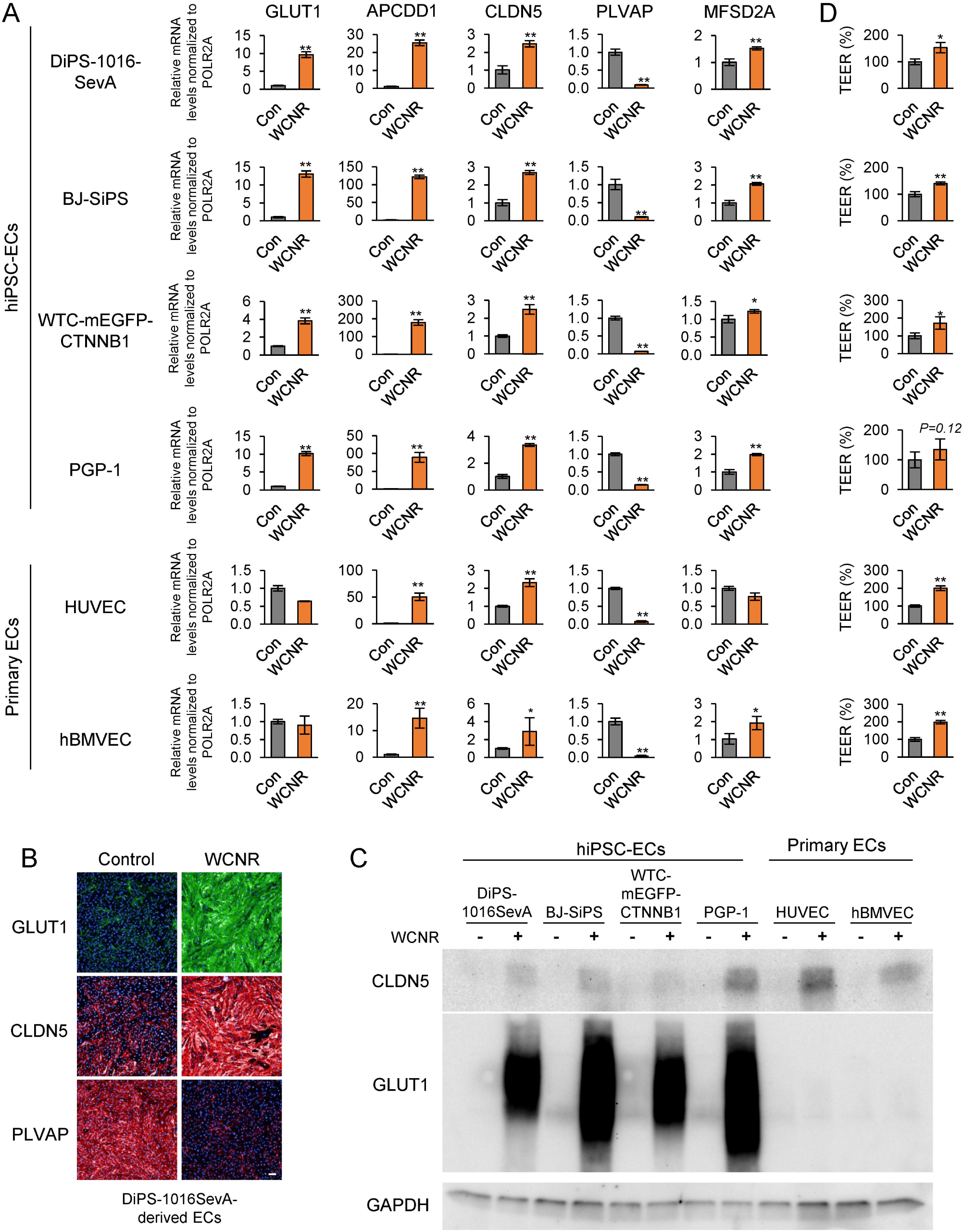
Combinatorial treatment of WNT3A, CNTF, CNTFRa, and RepSox (WCNR) induces brain EC-like phenotypes in hiPSC-ECs. (A) Relative mRNA expression of GLUT1, APCDD1, CLDN5, PLVAP, and MFSD2A after 2 days of WCNR treatment in hiPSC-ECs and primary ECs. (B) Representative images of cells treated with WCNR (100 ng/ml of rhWNT3A, 25 ng/ml of CNTF, 50 ng/ml of CNTFRa, and 10 uM of RepSox) for 2 days. Scale bar, 100 um. (C) Western blot showing CLDN5 and GLUT1 protein expression in hiPSC-derived ECs treated with WCNR. GAPDH was used as loading control. (D) Transendothelial electrical resistance (TEER) of control and WCNR-treated hiPSC-ECs after 4 days of treatment. Data are represented as mean ± standard deviation.

To examine the broader impact of WCNR on the brain EC-related transcriptional signature, we performed RNA sequencing. WCNR resulted in differential expression of a vast number of genes (Fig 5A). We analyzed a curated set of genes that are known to be expressed at least two-fold higher in brain ECs compared to peripheral ECs in mouse studies, as well as genes previously reported to be associated with BBB function(Cui et al., 2025; Hupe et al., 2017a; Munji et al., 2019; Sabbagh and Nathans, 2020b; Sabbagh et al., 2018) (Fig 5B and 5C). WCNR treatment caused expression changes in many of these genes, although not all. Among transporter genes, SLC2A1 (GLUT1), the xenobiotic transporter ABCG2, the retinol transporter STRA6, and the amino acid transporter SLC38A5 were increased. Canonical WNT downstream transcription factors LEF1 and TCF7, as well as the brain EC associated transcription factors ZIC3 and PPARD, were also induced. Tight junction related genes, including CLDN5, LSR, and ILDR2 were upregulated. In contrast, OCLN expression was reduced, which may partially explain the modest increase in TEER. Within categories related to metabolism, cell signaling, cytokine, or extracellular matrix (ECM), WCNR induced a portion of brain EC-enriched genes; however, approximately half of the surveyed genes were either unchanged or downregulated, indicating incomplete acquisition of a brain EC transcriptional profile (Fig 5C). We further examined genes involved in transcytosis and vesicle trafficking, processes that are suppressed in brain ECs, as well as pan-endothelial genes (Fig 5B). PLVAP was markedly downregulated by WCNR, consistent with qPCR data. In contrast, some other transcytosis or receptor-mediated endocytosis related genes, such as TFRC, did not display a consistent pattern of regulation. Among pan-EC genes, cell adhesion molecules like SELE, SELP, ICAM2 were reduced following WCNR treatment. Together, these data indicate that WCNR partially shifts hiPSC-derived ECs toward a brain EC-like transcriptional state, while additional cues likely are required to achieve complete brain EC differentiation as well as the acquisition of high electrical resistance tight junctions.

**Figure 5.**
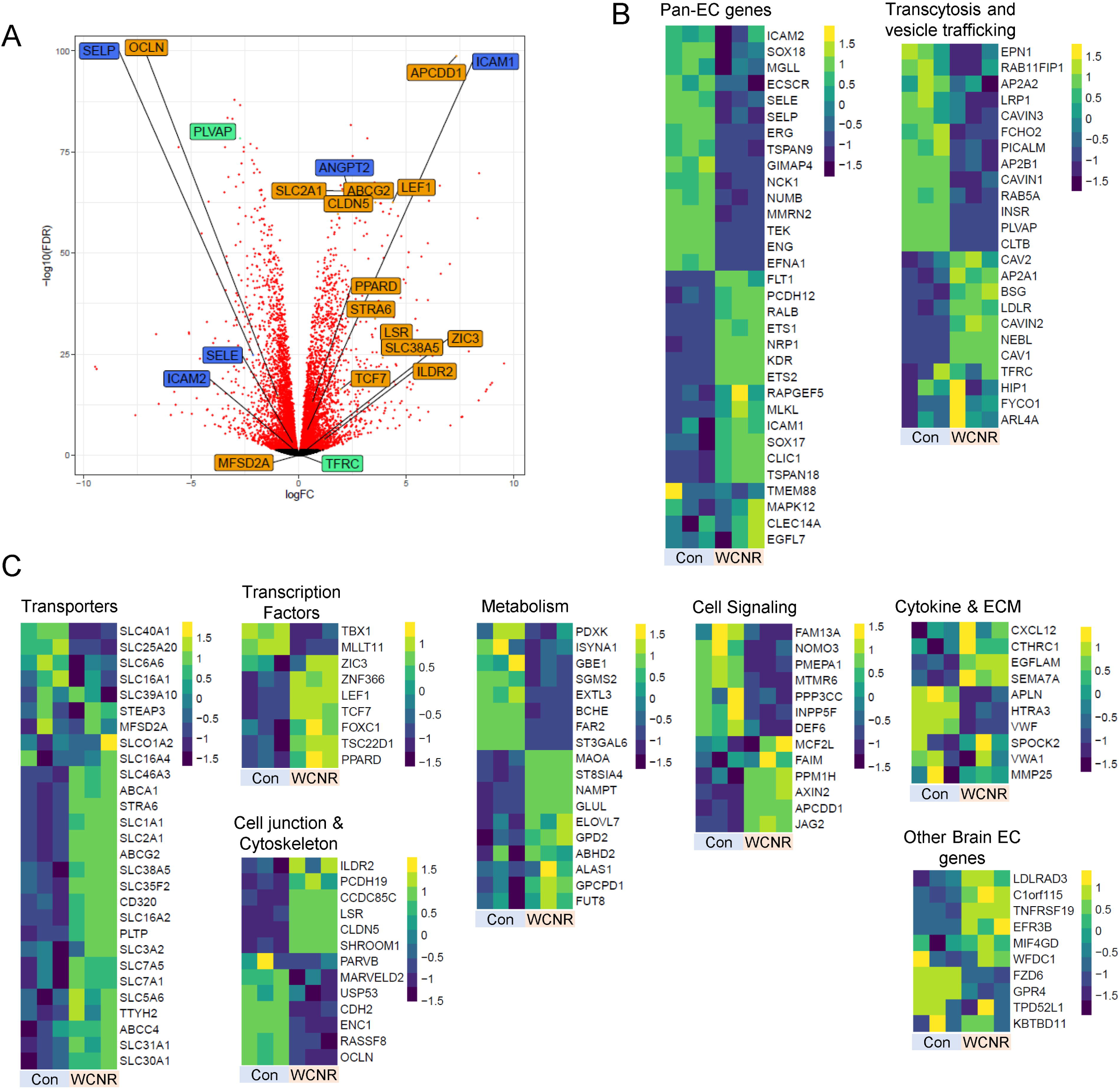
WCNR impacts broad BBB-associated genes in hiPSC-ECs. (A) Volcano plot showing differentially expressed genes in WCNR-treated hiPSC-ECs. (B) Heatmaps of pan-EC genes and transcytosis-related genes from RNA sequencing analysis comparing control and WCNR-treated cells. (C) Heatmaps of selected brain EC-enriched genes from RNA sequencing analysis comparing control and WCNR-treated cells.

### Inhibition of angiopoietin 2 signaling increases TEER

Through RNA sequencing, we discovered that angiopoietin-2 (ANGPT2), a secreted protein antagonist of the TEK receptor tyrosine kinase (Tie2)(Maisonpierre et al., 1997), was one of the genes induced by WCNR treatment (Fig 6A). ANGPT2 is a key regulator of vascular remodeling and is well known to increase endothelial permeability(Akwii et al., 2019; Brindle et al., 2006; Gurnik et al., 2016; Hakanpaa et al., 2015). When we measured the level of ANGPT2 in the conditioned medium of hiPSC-derived ECs, we found that it was extremely high (18,000-35,000 pg/ml) and increased further following WCNR treatment (Fig 6B). These levels greatly exceed those found in healthy human serum (1,000-4,000 pg/ml) and are in a similar range to those observed in patients suffering from a variety of traumatic injuries or during pregnancy, conditions associated with significant vascular remodeling (Hirokoshi et al., 2005; Van Hulle et al., 2024; Negrin and Hajdu, 2023).

**Figure 6.**
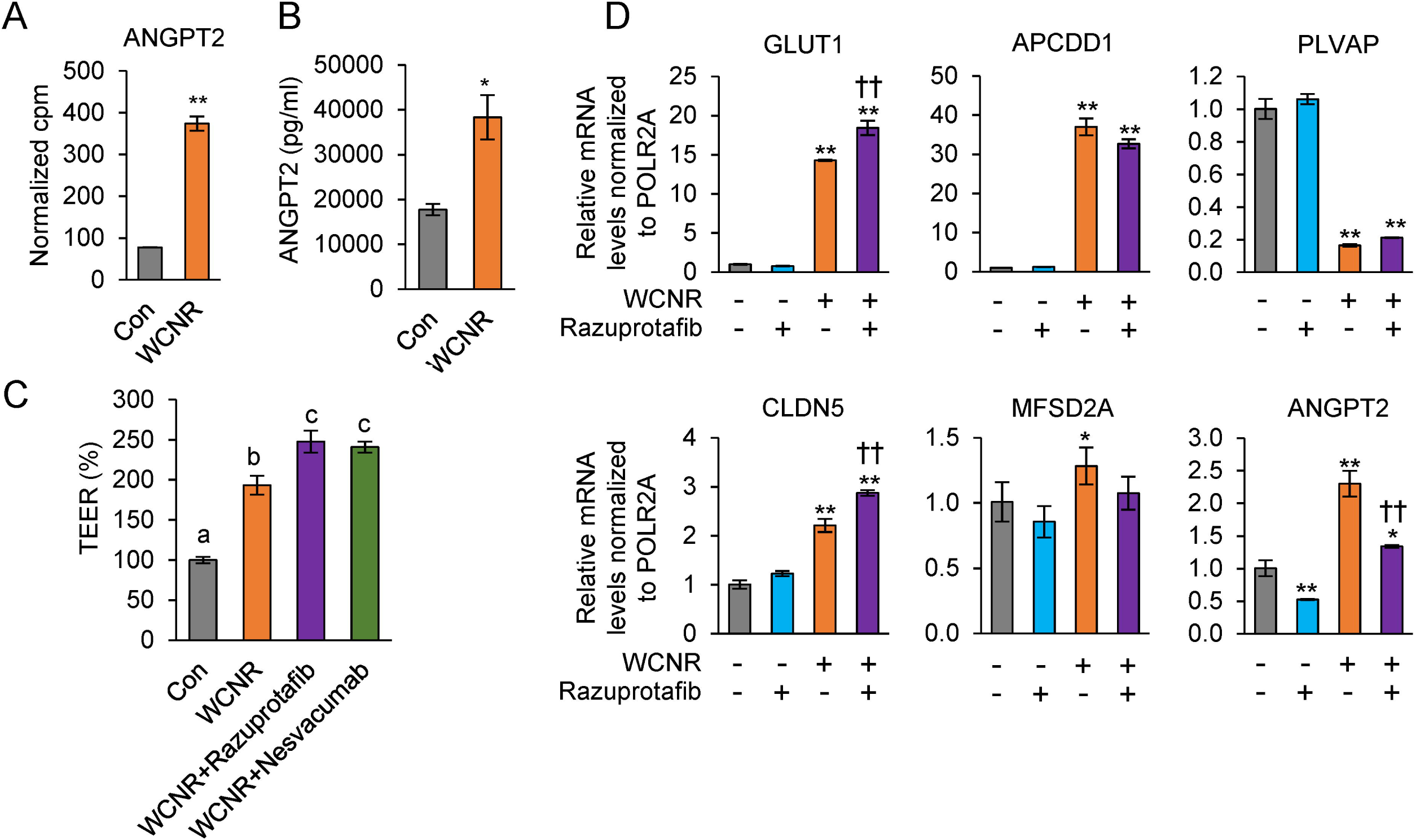
Inhibition of ANGPT2 signaling enhances barrier integrity in hiPSC-ECs. (A) Normalized counts-per-million (cpm) values from RNA sequencing of control and WCNR-treated hiPSC-derived ECs. WNT3A, CNTF, CNTFRa, and RepSox were treated for 2 days. **, FDR<0.01 (edgeR). (B) ELISA measurement of human ANGPT2 level in conditioned media from hiPSC-ECs. (C) TEER measurement of control, WCNR-treated, WCNR+razuprotafib-treated and WCNR+Nesvacumab-treated hiPSC-ECs after 4 days. Razuprotafib, 5 uM. Nesvascumab, 20 ug/ml. Different letters indicate significant differences between treatments (one-way ANOVA, Tukey’s HSD, p<0.05). (D) Relative mRNA expression of GLUT1, APCDD1, PLVAP, CLDN5, MFSD2A, and ANGPT2 in hiPSC-ECs treated with razuprotafib or WCNR. Data are represented as mean ± standard deviation. *, P<0.05 vs Control; **, P<0.01 vs Control; ††, P<0.01 vs WCNR, t-test.

To counteract the effects of elevated ANGPT2, we first incubated WCNR-treated hiPSC-ECs with razuprotafib (AKB-9778), a small molecule inhibitor of protein tyrosine phosphatase receptor type B (PTPRB; VE-PTP) that enhances TIE2 and reduces blood vessel permeability (Shen et al., 2014). Razuprotafib significantly increased TEER in WCNR-treated iPSC-ECs (Fig 6C). Similarly, treatment with Nesvacumab (REGN910), a humanized antibody that neutralizes ANGPT2, also increased TEER, further supporting the conclusion that excessive ANGPT2 limits barrier formation in cultured ECs. qPCR analysis indicated that razuprotafib alone did not significantly alter the expression of GLUT1, CLDN5, or PLVAP, although it reduced ANGPT2 expression (Fig 6D). However, co-treatment of razuprotafib and WCNR resulted in higher GLUT1 and CLDN5 expression than WCNR treatment alone, suggesting that Tie2 signaling may support the acquisition of brain endothelial characteristics in addition to enhancing barrier integrity. Collectively, these experiments demonstrate that individual signaling pathways regulate distinct components of the brain EC phenotype. No single treatment was sufficient to alter all of the selected brain EC–associated genes, whereas combinatorial pathway modulation produced broader and more pronounced changes toward a brain EC–like phenotype (Table 1).

**Table 1.** Regulation of Brain EC-associated Genes by WNT3A, CNTF, RepSox, and Razuprotafib.

| Genes | WNT3A | CNTF | RepSox | WCNR | Razuprotafib | WCNR+Razuprotafib |
| --- | --- | --- | --- | --- | --- | --- |
| GLUT1 | ↑↑ | ↑ | ↑ | ↑↑ | - | ↑↑↑ |
| PLVAP | ↓↓ | - | ↓ | ↓↓ | - | ↓↓ |
| CLDN5 | - | ↑ | ↑ | ↑↑ | - | ↑↑↑ |
| MFSD2A | - | - | - | ↑ | - | - |

↑, increased mRNA expression; ↓, decreased mRNA expression; -, no significant change. Additional arrows representing a stronger change in expression.

## Discussion

Our goal was to reproduce *in vitro* the normal developmental trajectory of ECs as they begin to vascularize the brain and acquire the specialized properties that define ECs that form the BBB. In doing so, our study provides valuable insight into several signaling pathways that regulate brain EC differentiation. During brain development, cell-cell interactions presumably provide a complete inductive environment whose molecular underpinnings are incompletely understood. *In vitro,* our approach indicates that simultaneous modulation of multiple pathways is required to induce brain EC-like phenotypes. Each pathway contributes to different aspects of brain EC differentiation. The canonical Wnt pathway strongly upregulates GLUT1 while downregulating PLVAP. STAT3 activation and TGFBR inhibition synergistically upregulate levels of the brain EC-specific tight junction protein CLDN5. Finally, inhibition of ANGPT2, which is present at supraphysiological levels in culture media, increased barrier strength by putatively stabilizing EC junctions. Together, these pathways partially reconstitute transcriptional and functional aspects of brain ECs derived from hiPSCs.

Wnt pathway activation by WNT3A robustly induced GLUT1 and suppressed PLVAP in hiPSC-derived ECs. WNT3A is a potent Wnt ligand capable of interacting with a broad spectrum of Frizzled receptors(Voloshanenko et al., 2017) and has been widely used to induce BBB-like properties in cultured ECs including immortalized brain ECs (Guérit et al., 2021; Laksitorini et al., 2019), primary brain ECs(Liebner et al., 2008), and iPSC-derived ECs(Praça et al., 2019). *In vivo*, Wnt7a and Wnt7b expression correlate strongly with β-Catenin activation in CNS vasculature, and double knockout of these ligands causes severe defects in brain vascular development (Daneman et al., 2009; Stenman et al., 2008). It is possible that the lack of responsiveness to recombinant WNT7A in our *in vitro* system is due to insufficient availability of appropriate receptor and co-receptors (GPR124, RECK, and LRP5/6) necessary for transmitting the WNT7A/WNT7B signal(Cho et al., 2017; Heiden et al., 2025; Vanhollebeke et al., 2015). Importantly, WNT3A induced GLUT1 to a degree comparable to CHIR99021 without massive nuclear localization of β-catenin, similar to a prior finding that β-catenin is highly active in embryonic brain ECs but much less active in postnatal stages(Liebner et al., 2008). This suggests that GLUT1 induction does not require sustained nuclear β-catenin, and that moderate activation of canonical Wnt signaling may be physiologically relevant. Furthermore, we found that primary human ECs like HUVECs and hBMVECs did not exhibit GLUT1 upregulation in response to WCNR treatment. This finding suggests that hiPSC-derived ECs may retain a more developmentally plastic state, allowing WNT signaling to induce GLUT1 expression.

Our data also show that STAT3 activation by CNTF contributes to CLDN5 induction in hiPSC-derived ECs. In the developing mouse brain, ECs express higher levels of Cntfr and Osmr compared to peripheral ECs (Hupe et al., 2017b), supporting our data that, among IL-6 family cytokines tested, CNTF and OSM, but not IL-6 or LIF, are capable of upregulating CLDN5. Although IL6ST (gp130)-STAT3 signaling can be triggered by multiple IL-6 family ligands including CNTF, OSM, IL-6, LIF, or cardiotrophin-1 (CTF1) depending on receptor availability, downstream effects of STAT3 in ECs are context-dependent. STAT3 activation by IL-6 or VEGF has been shown to increase vascular permeability (Wang et al., 2021b; Yun et al., 2017) whereas endothelial STAT3 has been reported to play a protective role in maintaining vascular integrity (Davis et al., 2022; Zhang et al., 2025). Together, our findings support a model in which gp130-mediated STAT3 activation promotes barrier formation through CLDN5 upregulation during brain EC differentiation.

The role of TGF-β signaling in vascular development and homeostasis is complex (Rattner et al., 2022). We found that inhibition of TGFBR1/ALK5 via RepSox strongly increased the expression of the tight junction protein CLDN5 while downregulating PLVAP. This is consistent with earlier reports(Roudnicky et al., 2020a; Watabe et al., 2003) and the well-established role of TGF-β in increasing endothelial permeability in culture(Behzadian et al., 2001; Goldberg et al., 2002; McMillin et al., 2015). TGF-β signaling can also drive an endothelial-mesenchymal transition (EndMT) in endothelial cells, a process associated with loss of tight junctions and increased plasticity (Ma et al., 2020). Thus, inhibition of TGF-b may enhance brain EC characteristics by preventing EndMT-like dedifferentiation in culture. Because ECs express TGF-β, and serum-containing medium introduces additional active and latent TGF-β (Danielpour et al., 1989; Oida and Weiner, 2010), basal TGFBR1 activity likely contributes to the default state of cultured ECs.

A key finding of our study is that hiPSC-derived ECs secrete a very high level of ANGPT2. Similar observations have been reported in RNA sequencing data of primary brain ECs(Sabbagh and Nathans, 2020a), where Angpt2 expression was significantly increased after isolation and culture, suggesting that the *in vitro* environment induces a stressed EC state. ANGPT2 is typically elevated under inflammatory or vascular injury conditions, and its overproduction likely contributes to junctional instability and barrier weakness. In addition, activation of Tie2 signaling through inhibition of VE-PTP has been shown to restore BBB integrity in aged mice (Todorov et al., 2026). Likewise, we found that the VE-PTP inhibitor Razuprotafib increased TEER in WCNR-treated ECs. Together, our study demonstrates that excess ANGPT2 is a substantial factor limiting barrier integrity in cultured hiPSC-derived ECs, and that restoring Tie2 activation can stabilize endothelial barrier function. It is plausible that additional stress-associated factors, aside from ANGPT2, also impede the acquisition of brain EC characteristics *in vitro*. Therefore, recreating a more physiologically relevant microenvironment including shear stress, ECM mechanics, and physiological oxygen tension may be necessary to achieve more faithful brain EC differentiation in vitro.

Among the four essential EC marker genes we focused on, MFSD2A was not induced by any of the pathways tested. Although β-Catenin has been shown to activate MFSD2A promoter activity in HEK293 cells (Wang et al., 2020), Wnt activation alone was not sufficient to induce MFSD2A expression in hiPSC-derived ECs. It is likely that MFSD2A requires additional unknown inductive cues. Alternatively, regulatory elements in the MFSD2A promoter region may be epigenetically repressed in cultured ECs. The *in vitro* culture environment, frequent dissociation and lack of supporting cells, may actively promote dedifferentiation of brain ECs, as seen in primary brain ECs that rapidly lose BBB properties (Sabbagh and Nathans, 2020a), thereby preventing ECs from acquiring brain EC characteristics. Understanding how cultured brain ECs lose and potentially regain their brain EC-specific identity may provide a strategy to further enhance brain EC differentiation from hiPSCs.

Taken together, this study characterizes the roles of multiple signaling pathways in the acquisition of key brain EC markers using hiPSC-derived ECs. Our findings demonstrate that the *in vitro* culture environment imposes limitations on brain EC differentiation, suggesting that overcoming culture-induced constraints, along with introducing the appropriate inductive cues, may be necessary to achieve more complete brain EC differentiation. These insights offer a foundation for improving brain EC differentiation strategies and advancing the development of more physiologically-relevant human BBB models. Our data may also have significant applications in elucidating how the BBB is regulated *in vivo* during development and in situations in which permeability is altered. The may also suggest means of regulating BBB permeability for the purpose of enhancing delivery of therapeutics to brain.

## Resource availability

### Lead contact

Requests for further information and resources should be directed to and will be fulfilled by the lead contact, Dr. Lee L. Rubin

### Materials availability

This study did not generate any new reagents.

### Data and code availability

RNA-seq data will be deposited upon acceptance to the Gene Expression Omnibus (GEO) managed by the National Center for Biotechnology Information

## Acknowledgement

Research reported in this publication was supported by Simons Foundation Collaboration on Plasticity in the Aging Brain (SCPAB); the Harvard Stem Cell Institute; a gift from the Vranos Family Foundation; a gift from the Edwin E McAmis Fund for Neuroscience; the National Institute of Neurological Disorders and Stroke (NINDS) of the National Institutes of Health under Award Number R01NS117407, and the National Institute on Aging (NIA) of the National Institutes of Health under Award Number R01AG072086. The total federal funds received for the NINDS award were $2,957,500 for a 5-year period, and NIA award for $3,129,600 for a 4-year, 8.5 months period, including direct and indirect costs. The content is solely the responsibility of the authors and does not necessarily represent the official views of the National Institutes of Health.

## Author contributions

J.H.L. and L.L.R. conceptualized the study and designed experiments. J.H.L., E.S.O., and J.Y.L. performed the experiments and data analysis. K.M.H. analyzed next-generation sequencing data. J.H.L. and L.L.R. wrote the manuscript. J.H.L., E.S.O., J.Y.L., K.M.H, and L.L.R. edited and revised the manuscript.

## Declaration of interests

L.L.R. is a founder of Vesalius Therapeutics, Valid Therapeutics, and iOrganBio, a member of their scientific advisory boards and a private equity shareholder. He is also a scientific advisory board member of ProjenX, Corsalex, and Jocasta Neurosciences. All are interested in formulating approaches intended to treat diseases of the nervous system and other tissues. None of these companies provided any financial support for the work in this paper. J.H.L. and L.L.R. are inventors on patent application PCT/US2023/23906, “Methods of Making Brain Endothelial Cells and Uses Therefor,” assigned to the President and Fellows of Harvard College. All other authors declare no competing interests.

## Materials and methods

### Human iPSC culture

All human iPSCs were cultured at 37°C in Stemflex media (Life Technologies A3349401) in plates coated with Geltrex LDEV-Free Reduced Growth Factor Basement Membrane Matrix (Life Technologies A1413202). Culture medium was replaced every 2-3 days and cells were passaged at approximately 80% confluency with 0.5 mM EDTA (Life Technologies, 15575020). The DiPS-1016SevA hiPSC (Harvard Stem Cell Institute, HVRDi007-A; RRID:CVCL_UK18) was mainly used for EC differentiation, RNA analysis, protein analysis, and imaging. CTNNB1::GFP iPSC line (Allen Cell, AICS-0058-067iPSC), obtained from Corielle Institute for Medical Research, was used for imaging β-Catenin and immunostaining. All cell cultures were maintained under sterile conditions and routinely tested for mycoplasma contamination.

### EC differentiation

Differentiation of endothelial cells was done as previously reported(Patsch et al., 2015b) with slight modification. Briefly, hiPSCs were dissociated into single-cell state with Accutase (StemCell Technologies Inc. 07920) and 1.8 x 10^6^ cells were plated into Geltrex-coated 10 cm dish in Stemflex with 10 uM ROCK inhibitor Y-27632. After 24 hours, cells were treated with 1:1 mixture of Neurobasal medium and DMEM/F-12 GlutaMAX supplemented with N2, B27 (all Life Technologies), 8 uM of Chir99021 (Cayman Chemicals), and 25 ng/ml of BMP4 (Peprotech) for 3 days. Then, the medium was replaced with StemPro-34 with 200 ng/ml of VEGF (Peprotech) and 2 uM of forskolin (Selleckchem). After 2 days, differentiated cells were detached using Accutase for endothelial cell purification. MACS was performed using CD144 microbeads and LS columns (Miltenyi Biotech) according to the manufacturer’s protocol. For purified EC culture, fibronectin (Corning 356008) was first reconstituted in sterile water to make 1 mg/ml stock solution. Then, the fibronectin solution was diluted in Phosphate Buffered Saline (PBS; Corning 21-040-CM) to 5 ug/ml and added to culture dishes to cover the entire surface. After 1 hour incubation at room temperature, fibronectin solution was removed and purified hiPSC-ECs were plated EGM-2 (Lonza CC-3162) at 2 x 10^5^ to 3 x 10^5^ cells/cm^2^ density in fibronectin coated plates. EGM-2 media was replaced every 2 days until cells reach confluency. Upon confluency, hiPSC-ECs were dissociated with TrypLE Express (Gibco 12605010) and plated into desired type of plates for the further experiment.

Unless otherwise specified, all reagents were administered to confluent, differentiated hiPSC-derived ECs maintained in EGM-2 medium. For WCNR treatment, confluent differentiated hiPSC-ECs were treated with recombinant human WNT3A (R&D systems, 5036-WN; 100 ng/ml), recombinant human CNTF (Miltenyi biotech, 130-138-928; 50 ng/ml), recombinant human CNTFRa (R&D systems, 303-CR; 25 ng/ml), and RepSox (Selleckchem, S7223; 10 uM) in EGM2 medium. The culture medium was refreshed every 2 days.

### Primary endothelial cell culture

For human primary EC culture, tissue culture plates were coated with 0.1% gelatin for 1 hour at room temperature. Human umbilical vein endothelial cells (HUVECs, Lonza) were maintained in EGM-2 medium (Lonza, CC-3162). Human brain microvascular endothelial cells (hBMVECs) (iXCells Biotechnologies) were maintained in Endothelial Cell Growth Medium (iXCells Biotechnologies, MD-0010) according to the manufacturer’s instructions. Primary ECs were used for experiments within five passages.

### RNA analysis

For mRNA level analysis, total RNA was extracted using TRIzol reagent (Life Technologies 15596026). For each sample, 500 ng of RNA per sample were used for cDNA synthesis using iSCRIPT cDNA synthesis kit (Bio-Rad Laboratories, 170-8891). Quantitative real-time PCR was performed using Fast SYBR Green Master Mix (Life technologies 4385614) on a Quant Studio 6 Flex Real-time PCR System (Applied Biosystems). Relative gene expression was calculated using 2^-ΔΔCt method. Statistical significance was calculated using ΔCt values, while 2^-ΔΔCt values were used for graphical representation. POLR2A or RPLP0 was used as a reference gene for normalization. For RNA sequencing, total RNA was isolated using the RNeasy micro kit (QIAGEN, 74004). RNA samples were submitted to Novogene (Sacramento, CA) for library preparation and sequencing. Briefly, poly(A)-enriched mRNA libraries were prepared and sequenced in a 150-bp paired-end configuration on an Illumina NovaSeq X Plus platform. Briefly, raw RNA-Seq reads were mapped onto the human genome using the STAR aligner version 2.7.9a against the Ensembl GRCh38 version 109 genome. Based on the counts table generated from STAR alignment, differential gene expression analysis was performed using edgeR version 3.40.2 with the quasi-likelihood F test to the TMM-normalized read counts, per contrast. TMM-normalized counts per million (CPM) were extracted for curated heatmaps and were log2 z-scored per gene, cut at +/- 2 using the pheatmap package version 1.0.12.

### Immunostaining

iPSC-ECs were cultured on fibronectin coated plates and were fixed with 4% paraformaldehyde (PFA) for 10 minutes at room temperature unless otherwise specified. For CLDN5 staining, cells were fixed with ice-cold methanol for 10 minutes on ice. Following fixation, cells were washed three times with PBS. PFA-fixed cells were permeabilized using 0.1% Triton X-100 in PBS for 10 minutes at room temperature. Cells were then incubated with blocking buffer (4% donkey serum, 0.1% Tween 20 in PBS) for 1 hour at room temperature. After blocking, samples were incubated with primary antibodies diluted in blocking buffer. The following primary antibodies were used: anti-PECAM1 (R&D Systems, AF806) anti-CDH5 (R&D Systems, AF938), anti-PLVAP (Abcam, ab81719), anti-GLUT1(Abcam, ab115730), anti-CLDN5 (Life Technologies, 35-2500), anti-ZO1 antibody (Life Technologies, 33-9100). The following day, samples were washed three times with PBS, incubated for 1 hour at room temperature with appropriate secondary antibodies and Hoechst (Invitrogen, H3569) in blocking buffer. After three additional PBS washes, images were acquired using the ImageXpress® Micro Confocal High-Content Imaging System (Molecular Devices).

### Western blotting

Total protein was extracted using RIPA buffer (Thermo, 89901) supplemented with protease and protein phosphatase inhibitor cocktail (Thermo, 78429 and 78426). Cells were lysed with RIPA buffer for 30 minutes on ice and centrifuged to collect supernatant. Protein quantification was determined using BCA protein assay kit (Thermo, 23227). Equal amounts of protein were mixed with LDS sample buffer (Invitrogen, NP0007) and incubated at 70°C for 10 minutes prior to separation by SDS-PAGE using Criterion TGX precast gels (Bio-Rad, 567-1124). Proteins were transferred to PVDF membranes using the Trans-Blot Turbo Transfer System (Bio-Rad). Membranes were blocked with 5% non-fat milk or bovine serum albumin (BSA) in TBS-T for 1 hour at room temperature, followed by incubation with primary antibodies overnight at 4°C. The following primary antibodies were used: anti-phospho-STAT3 (Y705) (Cell Signaling Technology, 9145), anti-STAT3 (Cell Signaling Technology, 9139), anti-β-tubulin (Abcam, ab6046), anti-CLDN5 (Life Technologies, 35-2500), anti-GAPDH (Abcam, ab9485), and anti-GLUT1 (Abcam, ab115730). The following day, membranes were washed three times with TBS-T (10 minutes each) and incubated for 1 hour at room temperature with HRP-conjugated secondary antibodies (anti-Rabbit IgG HRP, Life Technologies 31460; anti-Mouse IgG HRP, Life Technologies 31430) diluted in 1:10,000 in 5% non-fat. After three additional washes with TBS-T, chemiluminescent signal was detected using SuperSignal West Dura Extended Duration Substrate (Thermo, 37071) or SuperSignal™ West Femto Maximum Sensitivity Substrate (Thermo, 34095) and Biorad Chemidoc imaging system.

### Transendothelial electrical resistance (TEER) measurement

Differentiated ECs were seeded onto geltrex-coated transwells (Corning, 3460) at a density of 2 x 10^5^ cells per 12 mm diameter transwell insert. After 2 days, cells were treated with WCNR, razuprotafib, or nesvacumab as indicated. TEER was measured 2 or 4 days after the treatment. Prior to measurement, cultures were equilibrated to room temperature for 30 minutes. Electrical resistance was measured using Millicell® ERS-2 Voltohmmeter (Millipore, MERS00002) and EndOhm chamber (World Precision Instrument, ENDOHM-12G). Background resistance measured from geltrex-coated, cell-free transwell inserts was subtracted from each measurement. TEER values were expressed relative to the corresponding control condition.

### Statistical analysis

Unless otherwise stated, statistical significance was assessed using a two-tailed Student’s t-test. For comparisons involving multiple treatment groups, one-way ANOVA followed by Tukey’s post hoc test (Tukey’s HSD) was performed. A value of P < 0.05 was considered statistically significant compared to the control group. *, P<0.05 vs. control; **, P<0.01 vs. control.

## Supplemental Information

Document S1. Figures S1-S6

