## Supplemental File for "Combinatorial Modulation of Wnt, STAT3, TGF-β, and Tie2 Pathways Drives Brain Endothelial Cell–Like Differentiation from hiPSCs"

Document S1. Figures S1-S6

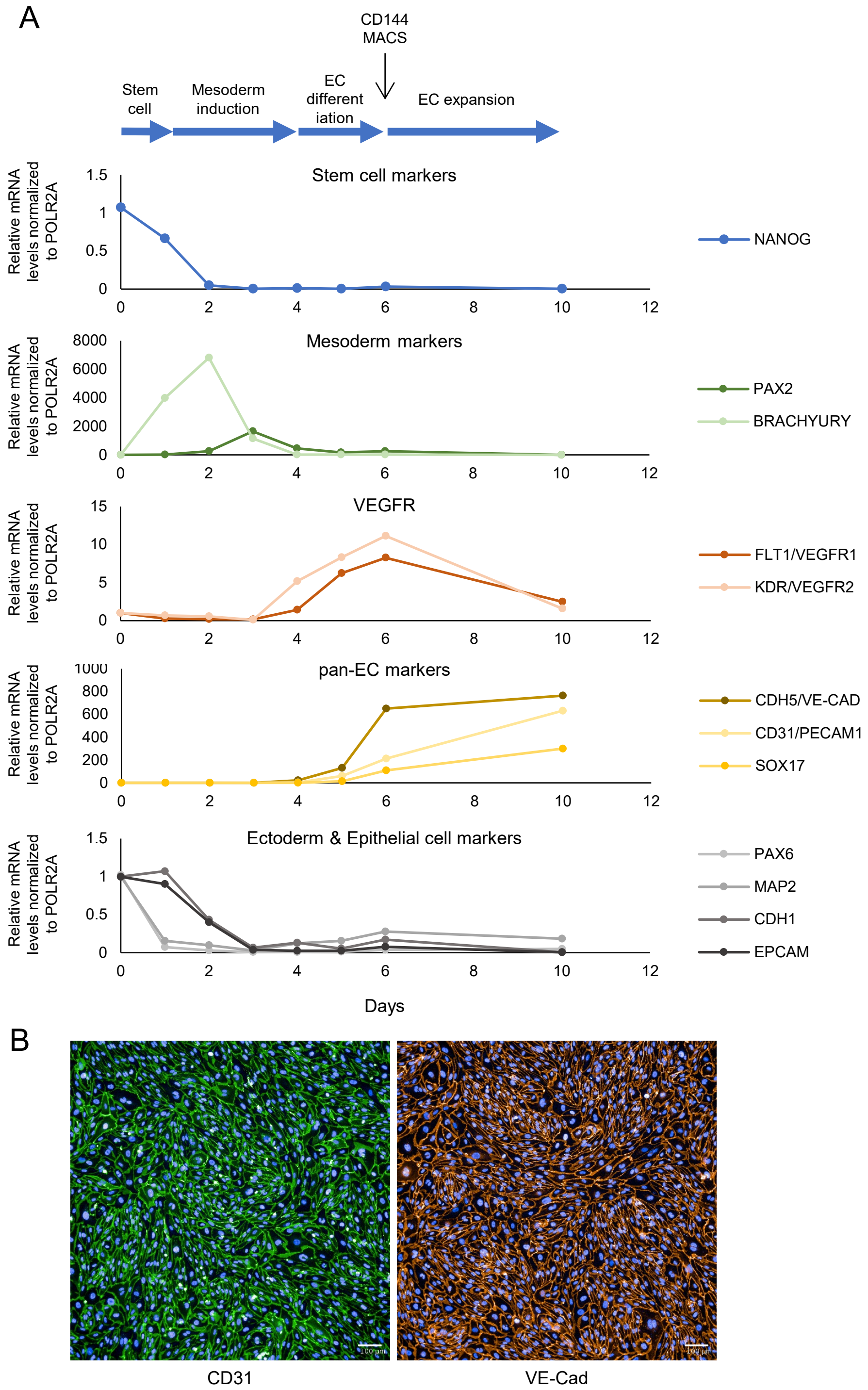


**Supplemental Figure S1. EC differentiation protocol generates homogeneous CD31- and CD144-positive cells.**

(A) mRNA expression of pluripotency, mesoderm, EC, and epithelial marker genes during differentiation from hiPSCs. (B) Immunofluorescence images of EC markers CD31(PECAM1) and CDH5 (VE-cadherin; CD144) following MACS. Scale bar, 100 um.

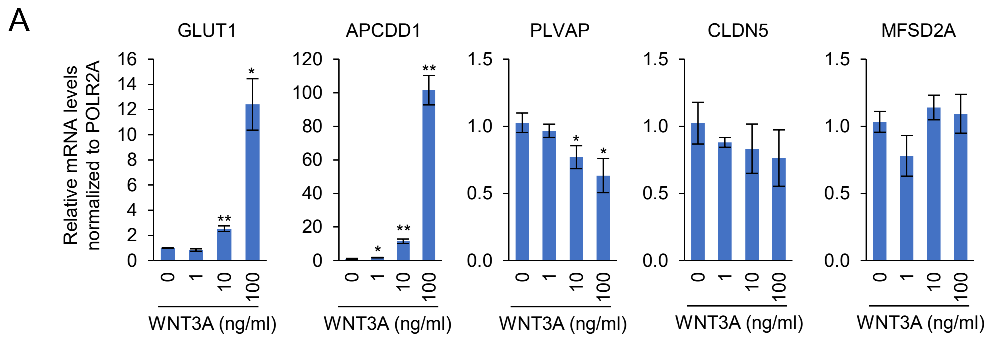


**Supplemental Figure S2. WNT3A induces GLUT1 and downregulates PLVAP.**
(A) mRNA expression of brain EC-associated genes in hiPSC-derived ECs after 2 days of treatment with varying concentrations of WNT3A. *, $P<0.05$ vs. control; **, $P<0.01$ vs. control.

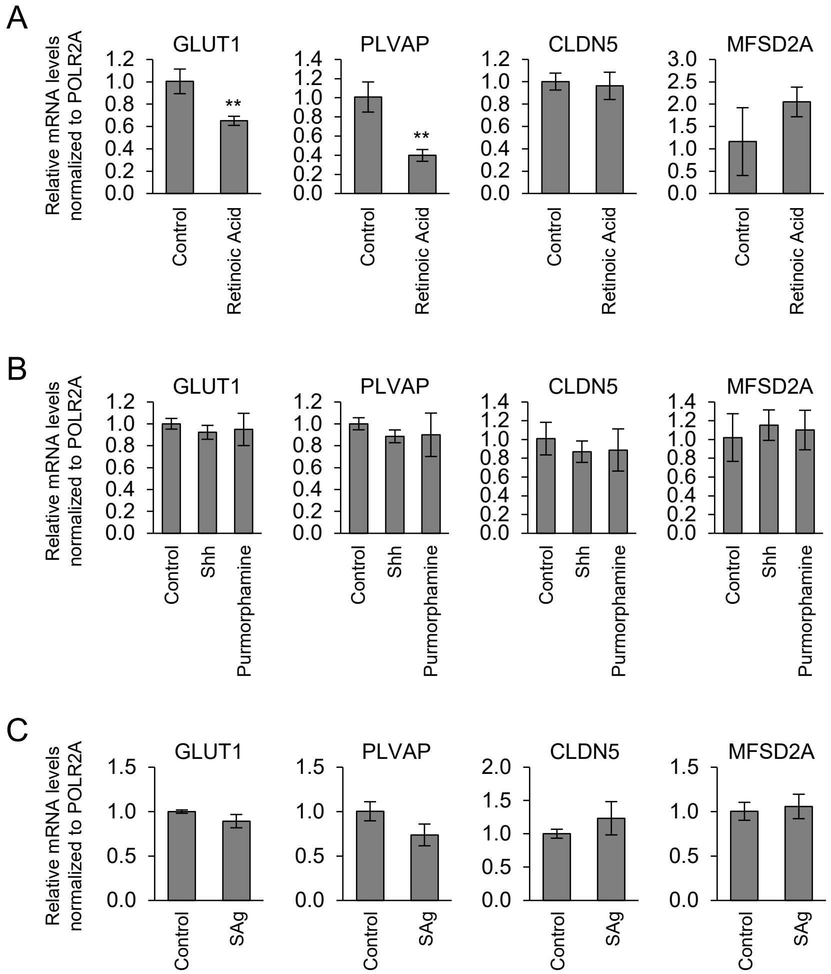


**Supplemental Figure S3. Retinoic acid and Hedgehog signaling do not promote brain EC differentiation.**

(A-C) mRNA expression of brain EC-associated genes in hiPSC-derived ECs after 2 days of treatment with retinoic acid (1 uM), recombinant human Sonic Hedgehog (Shh) (100 ng/ml; Peprotech #100-45), purmorphamine (1 uM; ReproCELL #04-0009), or Smoothened agonist (SAg) (1 uM; MilliporeSigma #566660). **, P<0.01 vs. control.

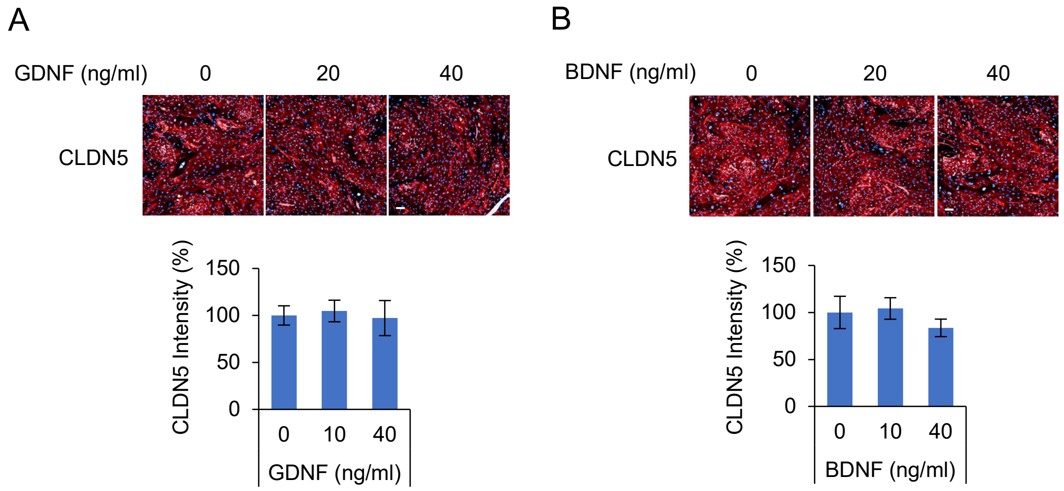


**Supplemental Figure S4. GDNF and BDNF do not increase CLDN5 expression in hiPSC-derived ECs.**

(A and B) Immunostaining and quantification of CLDN5 in iPSC-ECs treated with GDNF (A) or BDNF (B). GDNF and BDNF were treated at 0, 20, and 40 ng/ml in EGM2 media for 2 days. Scale bars, 100 um.

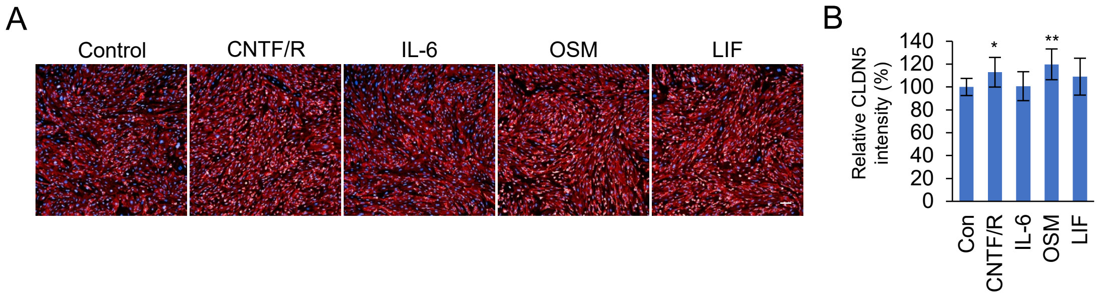


**Supplemental Figure S5. CNTF and Oncostatin M (OSM) induce CLDN5 expression in hiPSC-derived ECs.**

(A) CLDN5 Immunostaining in hiPSC-derived ECs treated with IL-6 family cytokines. (B) Quantification of CLDN5 intensity. Treatment concentrations: CNTF, 25 ng/ml; CNTFRα, 50 ng/ml; IL-6, 100 ng/ml; OSM, 100 ng/ml; LIF, 100 ng/ml. CNTF/R, CNTF and CNTFRα. Scale bars, 100 um.

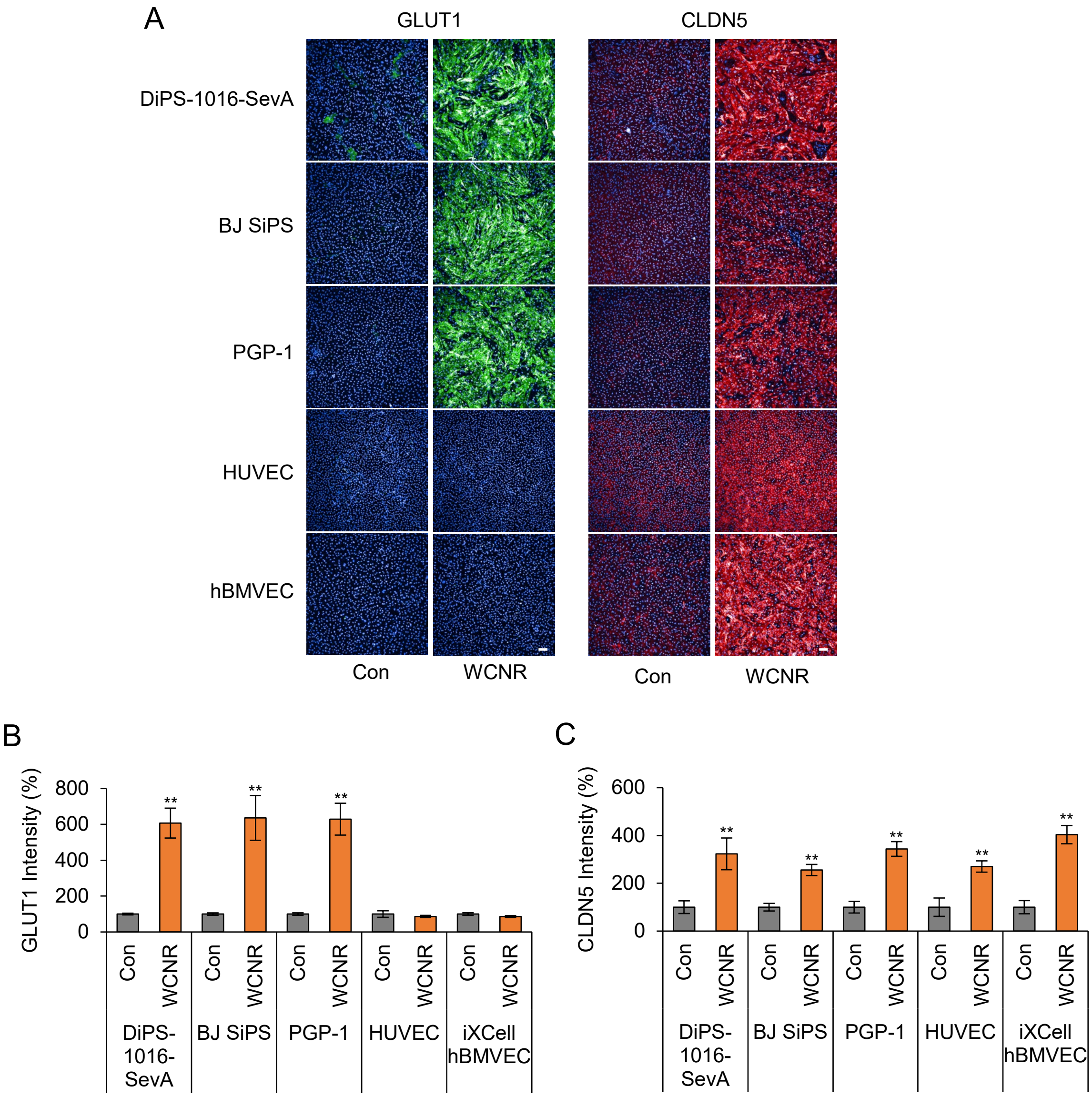


**Supplemental Figure S6. WCNR increased GLUT1 and CLDN5 in iPSC-ECs.**
(A) GLUT1 and CLDN5 immunostaining in multiple hiPSC-ECs lines (DiPS-1016-SevA, BJ SiPS, and PGP-1) and primary ECs (HUVEC, hBMVEC). WNT3A, CNTF, CNTFRα, and RepSox were treated for 2 days. (B and C) Quantification of GLUT1 and CLDN5 intensity, respectively. **, $P<0.01$ vs control (t-test).
